# Quantum and Quantum-inspired Methods for *de novo* Discovery of Altered Cancer Pathways

**DOI:** 10.1101/845719

**Authors:** Hedayat Alghassi, Raouf Dridi, A. Gordon Robertson, Sridhar Tayur

## Abstract

The combinatorial calculations for *de novo* discovery of altered pathways in cancer cohorts involve both coverage (i.e. recurrence) and mutual exclusivity, and solving mutual exclusivity problems is NP-hard for classical computers. Advances in quantum computing systems and our classical, quantum-inspired algorithm GAMA (Graver Augmented Multi-seed Algorithm) motivated us to revisit methods for identifying altered pathways. Using different types of quantum and classical algorithms, we report novel formulations of the problem that are tailored to these new computational models. Our formulations require fewer binary variables than available methods, and offer a tuning parameter that allows a trade-off between coverage and exclusivity; varying this parameter returns a variety of solutions. We illustrate our formulations and methods with TCGA mutation data for Acute Myeloid Leukemia (AML). Both the D-Wave quantum annealing solver and the classical GAMA solver returned altered pathways that are known to be important in AML, with different tuning parameter values returning alternative altered pathways. Our reduced-variable algorithm and QUBO problem formulations demonstrate how quantum annealing-based binary optimization solvers can be used now in cancer genomics.

## 1 Introduction

Identifying altered cancer pathways *de novo* from mutation co-occurrence and mutual exclusivity is NP-hard, primarily because of challenges in addressing mutual exclusivity (Karp, 1972). See, for example (Ciriello et al., 2012; Vandin et al., 2012; Leiserson et al., 2013; Zhao et al., 2012; Szczurek and Beerenwinkel, 2014; Leiserson et al., 2015), and reviews in (Deng et al., 2019; Dimitrakopoulos and Beerenwinkel, 2017; Cheng et al., 2016).

In parallel with advances in cancer genomics, there have been advances in computational models, notably in Adiabatic Quantum Computing (AQC), where the state of the art is a 2000-qubit machine from D-Wave Systems (Burnaby, Canada) (see Appendix C). While quantum computers will, in principle, speed up optimizations, they cannot yet handle the size of problems in cancer genomics, and this limitation will likely persist for some time (Preskill, 2018).

Recently, we reported the Graver Augmented Multi-seed Algorithm (GAMA) (Alghassi et al., 2019a), a classical optimization algorithm that was inspired by our work with quantum computing methods (Alghassi et al., 2019b). We have demonstrated that, for problem formulations that have a certain structure, GAMA’s optimization performance is superior to that of general purpose, commercial, classical solvers. As well, GAMA offers a spectrum of solutions that include all degenerate global solutions, and all suboptimal solutions.

In the work reported here, we revisit the problem of discovering altered cancer pathways from a mutated-gene-patient matrix, using novel formulations that are tailored to be solvable by the D-Wave D2000Q quantum computer, and, with a classical computer, by GAMA. Our problem modelling approach is new. We model the alteration matrix as a hypergraph and map it to its primal graph, and finally to the graph Laplacian. Our new problem formulations require a number of binary variables consisting of the total number of altered genes, rather than the sum of altered genes plus patients, which is required by *de novo* methods in the literature; thus, our problem formulation (used in QUBO or GAMA form) is more efficient than existing methods. Further, the levels of mutual exclusivity within and across cancer pathways vary; thus, rather than model exclusivity as a hard requirement, our formulations allow some degree of non-exclusivity, and we parameterize co-occurrence (coverage) and mutual exclusivity via a tuning parameter that allows us to vary their relative weights.

We illustrate our quantum and classical formulations for *de novo* identification of altered cancer pathways using mutation data from the TCGA acute myeloid leukemia (AML) study (Cancer Genome Atlas Research Network, 2013), and compare our solutions to those reported in that work.

## 2 Methods

### 2.1 Hypergraph Modeling: Gene as vertex, Patient as hyperedge

The alteration matrix *B* can be modeled as the incidence matrix of a hypergraph (Wikipedia, 2019) *H_g_* = (*V*_*g*_, *E*_*p*_), in which each vertex *v*_*i*_ ∈ *V*_*g*_, *i* = 1, 2, … , *n* represents a gene *v*_*i*_ ≡ *g*_*i*_, and each patient *P*_*i*_ represents a hyperedge *e*_*i*_ ∈ *E*_*p*_, *i* = 1, 2, … , *m*.

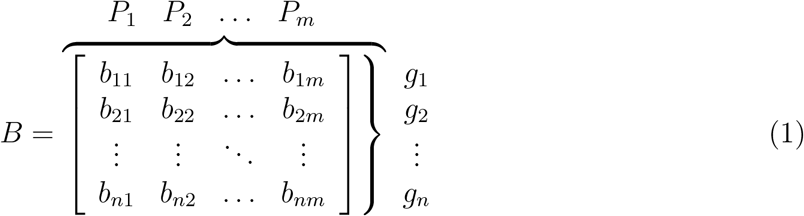

**Figure 1:**
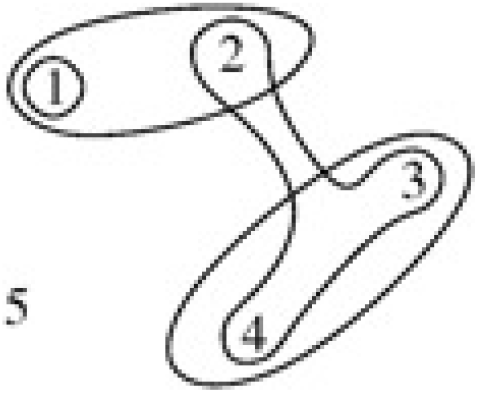
The hypergraph based modelling vertex = gene, hyperedge = patient.

Each column of *B* that represents a specific patient’s mutation list is a hyperedge. The primal^1^ graph (*G*) of a hypergraph (*H*) is a graph with the same vertices of the hypergraph and edges between all pairs of vertices contained in the same hyperedge.

The primal graph (*G*) of the hypergraph (*H*) can be found using the incidence matrix of the hypergraph *B*(*H*):

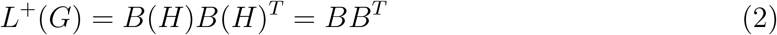

The matrix *L*^+^(*G*) = *D*(*G*) + *A*_*w*_(*G*), called the (positive) Laplacian matrix of *G*, is an *n* × *n* symmetric and positive definite matrix. The weighted adjacency matrix *A*_*w*_ = [*a*_*w*_(*i*, *j*)]^*n*×*n*^ is a symmetric and zero diagonal matrix in which *a*_*w*_(*i*, *j*) counts the number of patients that have gene pairs (*g*_*i*_, *g*_*j*_) mutated. Also *D* = *diag* {*d*_1_, *d*_2_, … , *d*_*n*_} is the degree matrix, in which *d_i_* is the degree of vertex *v*_*i*_ ≡ *g*_*i*_ in the primal graph, counting the number of patients that have gene *g*_*i*_ mutated.

Using a threshold value, the weighted edges in *A*_*w*_ become unweighted. The adjacency matrix of the unweighted graph is

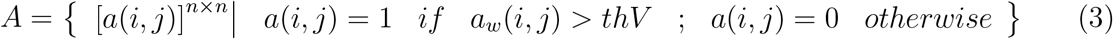

The *thV* is a threshold value that designates the minimum number of patients needed to have a mutation for a gene pair. This builds an unweighted edge between vertices (genes).

Two combinatorial criteria are important in the *de novo* discovery of cancer driver-gene pathways:

1. **Coverage:** The more cancer patients are affected by a mutated gene, the more likely it is that the gene is a driver gene.
2. **Exclusivity:** If a patient has a mutated driver gene in a pathway, the patient has no other mutation in other driver genes in the same pathway. Due to noise in signals, sample impurity, and other issues, the concept of exclusivity has been observed to be violated sporadically.

An independent (or stable) set of a graph is a subset of vertices of the graph whose elements are pairwise nonadjacent. The independent set with maximum cardinality is called the maximum independent set. If, instead of simple cardinality, each node is assigned a weight, the solution with the maximum total weight of vertices is called the *maximum weighted independent set*. The definition of independent sets in a hypergraph is as follows: given the hypergraph *H* = (*V*, *E*), find the maximum subset of *V* such that the vertex-induced subgraph on it does not contain a hyperedge.

For a mutation matrix *B*, the maximum mutual exclusivity of groups of genes is equivalent to the maximum independent set of the primal graph of the hypergraph in which genes are nodes and patient groups are hyperedges.

To satisfy both exclusiveness and maximum coverage, we need to find all independent sets of the primal graph and maximize over its nodal sum degrees. The problem has been transformed into finding the maximum weighted independent set of the primal graph of *H*_*g*_.

### 2.2 Formulation 1: Tailored for a Quantum Annealing Solver

#### 2.2.1 Single-Pathway QUBO problem (Formulation 1.1)

Quantum annealing-based processors in general, and the D-Wave D2000Q processor in particular, solve quadratic unconstrained binary optimization (QUBO) problems. In this framing, the main optimization term and constraints should be added together to form a single binary optimization problem with quadratic (i.e. two-body) interactions.

Recall that we want to find the independent set with the maximum coverage.

Assume **x** = [*x*_1_ *x*_2_ … *x*_*n*_]^*T*^ is a binary solution vector, indicating which nodes are in the independent set (pathway). The independence term can be written as 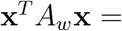 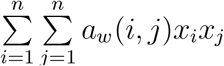, which becomes zero for any independent set. Also, the weighted coverage term can be written as 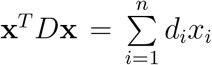; that is, the sum of node degrees (i.e. patients) should be maximized.

Thus, the QUBO formulation of the maximum coverage (weighted) independent set is

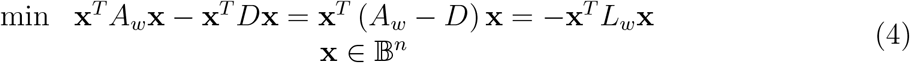

Here *A*_*w*_ is the adjacency matrix of the primal graph of the hypergraph, *D* is the degree matrix, and *L*_*w*_ = *D* − *A*_*w*_ is the (negative) Laplacian matrix.

To accommodate a trade-off between the independence term and the coverage term, we add a tuning parameter (*α*) so that we can flexibly balance the two terms:

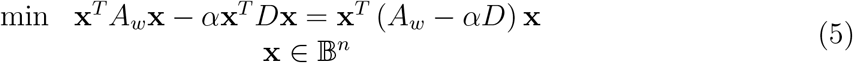

An important aspect of our formulation is that the term represented by **x**^*T*^ *D***x** is not exactly equal to the numerical coverage, but is a function that is monotonic with the exact coverage (i.e. their maxima are at the same argument vector). We discuss this further and compare our compact formulation with the maximum coverage exclusive submatrix formulation of (Vandin et al., 2012; Leiserson et al., 2013) in section 2.4 below. To find several distinct pathways, using the single pathway formulation, we iteratively solve (5) and remove the genes acquired from iteration *l* for iteration *l* + 1.

#### 2.2.2 Multiple-Pathway QUBO problem (Formulation 1.2)

To find *k* pathways simultaneously from one optimization run, we formulate the *k*-pathway QUBO. The binary vector **x**_*i*_ = [*x*_*i*1_ *x*_*i*2_ … *x*_*in*_]^*T*^, *i* = 1, 2, … , *k* is the solution vector, that indicates whether the vertex *v*_*j*_ belongs to the *i*^*th*^ pathway; when it does, *x*_*ij*_ is equal to 1, otherwise it is 0.

Let **X** = [**x**_1_ **x**_2_ … **x**_*k*_]^*T*^. The general QUBO problem formulation of the *k*-maximum weighted independent set is (Alghassi, 2015):

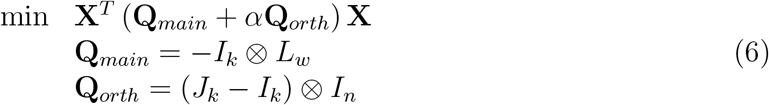

*I*_*k*_ and *I*_*n*_ are *k* × *k* and *n* × *n* identity matrices, and *J_k_* is the *k* × *k* matrix in which all entries are 1. (Recall that *α* is the orthogonality constraint balancing factor.) In the majority of cases, the cardinality of solutions is not known; however, if we know the size (cardinality) of the pathways, a cardinality constraint term^2^ can be added to the **Q**_*main*_ as second constraint in (6). The derivation of the tensor formulation of constraints is discussed in detail in (Alghassi, 2015).

### 2.3 Formulation 2: Tailored for the GAMA Solver

Here we modify the multiple-pathways model discussed in 2.2.2 for the Graver Augmented Multi-seed Algorithm (GAMA) solver (Alghassi et al., 2019a).

We can rewrite the formulation of (6) as

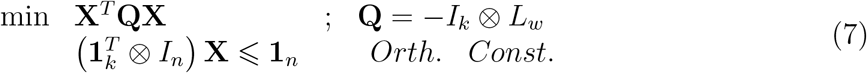

To transform the inequality to equality in orthogonality constraint, we add one extra binary slack variable for each constraint, thus resulting in (*k* + 1)*n* total binary variables. This is a nonlinear (quadratic) nonconvex integer problem that has the form of a Quadratic Semi Assignment Problem (QSAP). Similarly, a cardinality constraint^3^ can be added (if the cardinality is known), and the problem with both orthogonality and cardinality constraints becomes a Quadratic Assignment Problem (QAP).

Appendix B discusses briefly, and (Alghassi et al., 2019a) discusses in detail, systematic Graver basis extraction and the GAMA solver.

### 2.4 Comparison of Our Formulations with Existing Literature

We compare our formulations with the *maximum coverage exclusive submatrix* formulation presented in (Vandin et al., 2012). To show that our method requires fewer binary variables to model the same optimization problem, we first rewrite the maximum coverage exclusive submatrix in its original form.

Assume the set *M* contains genes with coverage and exclusivity requirements. Then, if Γ(*M*) represents the total number of patients for the gene set *M*, and Γ(*g*_*i*_) represents the number of patients with gene *g*_*i*_, mutual exclusivity holds when 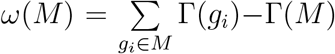
reaches its minimum value of zero.

By replacing the maximum coverage term with the minimum of its negative, the optimization model is as follows (Vandin et al., 2012; Leiserson et al., 2013):

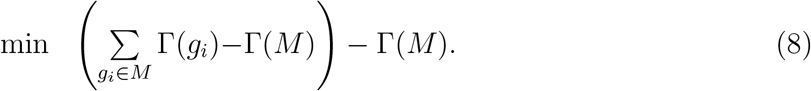

To implement this approach as a binary optimization problem, we need to have *n* binary variables **x** = [*x*_1_ *x*_2_ … *x*_*n*_]^*T*^ to identify genes that represent *M*, and another *m* binary *T* variables **y** = [*y*_1_ *y*_2_ … *y*_*m*_]^*T*^ to identify patients that have at least one mutated gene inside gene set *M*. Thus (8) can be written as

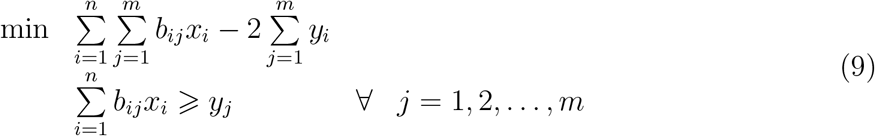

The above can be rewritten in matrix form:

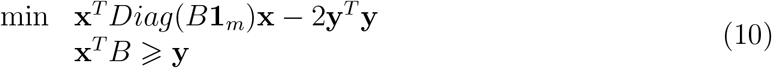

To rewrite the inequality constraint in (9) or (10) as an equality constraint, we need to introduce at least *m* extra integer slack variables^4^. Using the above approach requires at least *n* + 2*m* binary variables.

Our proposed optimization problem Formulation 1.1 (see equation (5)) also contains two terms:

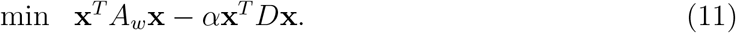

The first part models exclusiveness as finding the independent set of a weighted graph. The second part, which models coverage in reality, does not maximize the exact coverage (Γ(*M*)) but the sum of patients with genes listed 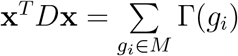. There is no need to maximize the real coverage, since Γ(*M*) and 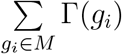 are monotone to each other and they achieve the same result, while total exclusiveness (independent set) is the solution. Our compact optimization model needs only *n* (number of genes) binary variables, which is more efficient, especially as the number of patients increases.

## 3 Results

### 3.1 Discovering Altered Pathways in AML using the D-Wave Solver

Using the gene-as-node, patient-as-hyperedge hypergraph model and its corresponding primal graph, we solved the QUBO problem formulation using the mutation data from the TCGA AML study (Cancer Genome Atlas Research Network, 2013). The mutation data was generated by genome and exome sequencing of samples from 200 patients. The gene set data was reduced to 33 genes, based on the method given in the publication’s supplementary document. The TCGA authors used Dendrix++ (Vandin et al., 2012; Leiserson et al., 2013, 2015) to report three disjoint pathways (groups of genes) that had the strongest patterns of mutual exclusivity. The size of the combinatorial optimization algorithm that we formulated and solved is equal to the total number of genes involved (*n* = 33), whereas in the optimization formulation presented in (Leiserson et al., 2013), on which the Dendrix++ approximate solution is based, the size of problem is equal to the number of genes plus the number of patients (*n* + *m* = 33 + 200 = 233). Using our sequential approach and the D-Wave quantum processor, we generated results similar to those reported for Dendrix++ (Appendix A).

### 3.2 Discovering Altered Pathways in AML using the GAMA Solver

We then used our reduced variable model for *k*-pathway extraction, formulated as a QSAP problem (sec. 2.3), and solved it with GAMA (Alghassi et al., 2019a) for the same AML mutation data (Cancer Genome Atlas Research Network, 2013). The size of integer optimization problem that we solved is (*k* + 1) times the total number of altered genes involved (*n* = 33), whereas, for the (Leiserson et al., 2013) model, the problem size would be *k* times (*n* + *m* = 33 + 200 = 233). Using the QSAP model and the GAMA solver we recovered largely the same results for *k* = 3 (Appendix A). For *k* = 6, three of the pathways are the main pathways that were previously reported, and have higher coverage, and the other three are suggested pathways that have lower coverage. We also show how, by changing the tuning parameter *α*, we can change the balance between the gene coverage and exclusivity among the *k* solutions.

## 4 Conclusions

We have developed novel hypergraph-based formulations of the *de novo* altered cancer pathway detection problem. Our formulations involve fewer variables, in that the number of binary variables in our method depends on the number of genes rather than the number of genes plus the number of patients. We have incorporated a tuning parameter between coverage and exclusivity (here, the independent set) to make alternative solutions flexibly available. We offer single-pathway and multiple-pathway QUBO formulations. We have also devised a (nonconvex and nonlinear) integer programming formulation for a multiple-pathway detection method based on a QSAP model. This formulation is suitable for the GAMA solver.

Using the D-Wave QUBO solver, we tested our QUBO formulations on TCGA AML mutation data, and recovered results similar to those reported in that work (see Appendix A). Similarly, we tested our multiple-pathway QSAP formulation with our GAMA solver.

In a companion study (Dridi et al., 2019), we expand the work reported here to address a topological perspective on pathways. We first formalize our tunable parameter concept using methods of algebraic topology. We then explore the shapes of cancer pathways; that is, we study how the various pathways are adjacent, and find interesting differences between AML and Glioblastoma Multiforme (GBM) cancer pathways.

Our reduced-variable algorithm and QUBO problem formulations are a gateway for using quantum annealing-based binary optimization solvers in cancer genomics in general. Here, we describe how to use such solvers to address *de novo* detection of altered pathways. The QSAP problem formulation of the algorithm, coupled with our quantum-inspired classical GAMA solver, fills a gap in small and noisy quantum solvers in the Noisy Intermediate-Scale Quantum (NISQ)(Preskill, 2018) era.

## Acknowledgements

The authors thank Sepideh M. Alamouti (Fusion Genomics Corporation, Burnaby, Canada) for helping us process genomic data. Work on QUBO formulations was initiated by the first author while working at 1QBit Technologies Inc. (Vancouver, Canada), and tested on an earlier version of D-Wave.

## A Appendix: Detailed Pathway Results on AML Data

## A.1 D-Wave Solver Results

Since in some practical cases of altered pathways the mutual exclusivity cannot be exact and some tolerance is permitted, the tuning parameter *α* is important for adjusting the balance between exclusivity and coverage. For the AML data, the results from our method are very similar to the results presented in (Cancer Genome Atlas Research Network, 2013) (page 2066), using *α* = 0.45.

Three main altered pathways obtained with our method are as follows:

- Pathway1 = ‘PML.RARA’, ‘MYH11.CBFB’, ‘RUNX1.RUNX1T1’, ‘NPM1’, ‘TP53’, ‘RUNX1’, ‘MLL-X fusions’, ‘MLL.PTD’, coverage = 141, coverage/gene = 17.63, indep = 8, measure = 2.20
- Pathway2 = ‘FLT3’, ‘Other Tyr kinases’, ‘Ser-Tyr kinases’, ‘KRAS/NRAS’, coverage = 112, coverage/gene = 28, indep = 14, measure = 2
- Pathway3 = ‘Other myeloid TFs’, ‘ASXL1’, ‘Other modifiers’, ‘Cohesin’, coverage = 68, coverage/gene = 17, indep = 0, measure = infinity

We have added an option to our code for filtering out the gene pairs for those cases in which the number of patients is lower than a threshold value (here 4). The resulting pathways are almost a match to the pathways presented in (Cancer Genome Atlas Research Network, 2013), except in the case of Pathway1, in which we have gene ‘MLL.PTD’ added instead of gene ‘CEBPA’. Checking the coverage and exclusivity shows that whereas ‘CEBPA’ has slightly better coverage, ‘MLL.PTD’ has better exclusivity.

Figure 2 depicts the above-mentioned results. In this graph representation, each gene is depicted as a node. The diameter of each node (gene) is proportional to the number of patients that carry the mutated gene. The edges and their thickness represent the number of patients who carry the mutated gene pairs linked with edge nodes.

**Figure 2:**
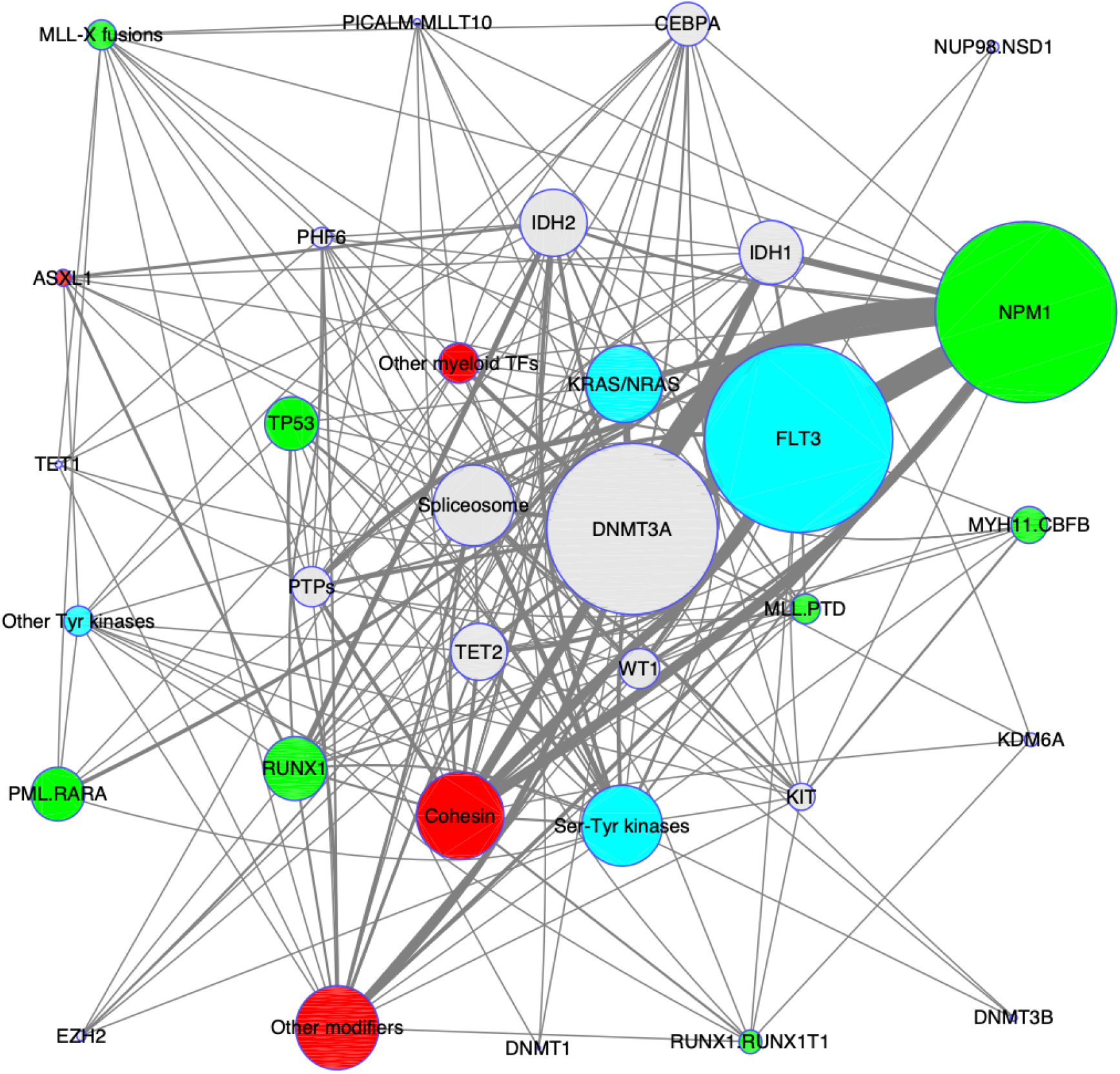
Graphical representation of AML data and three pathways identified with *α* = 0.45.

Increasing the value of *α* enhances the weight of coverage and relaxes the exclusivity principle, thus expanding the number of genes in each pathway. In the case of the AML data, increasing the value of *α* from 0.45 to 0.70 adds ‘CEBPA’ to the list of genes in Pathway1. This case is depicted in Figure 3.

**Figure 3:**
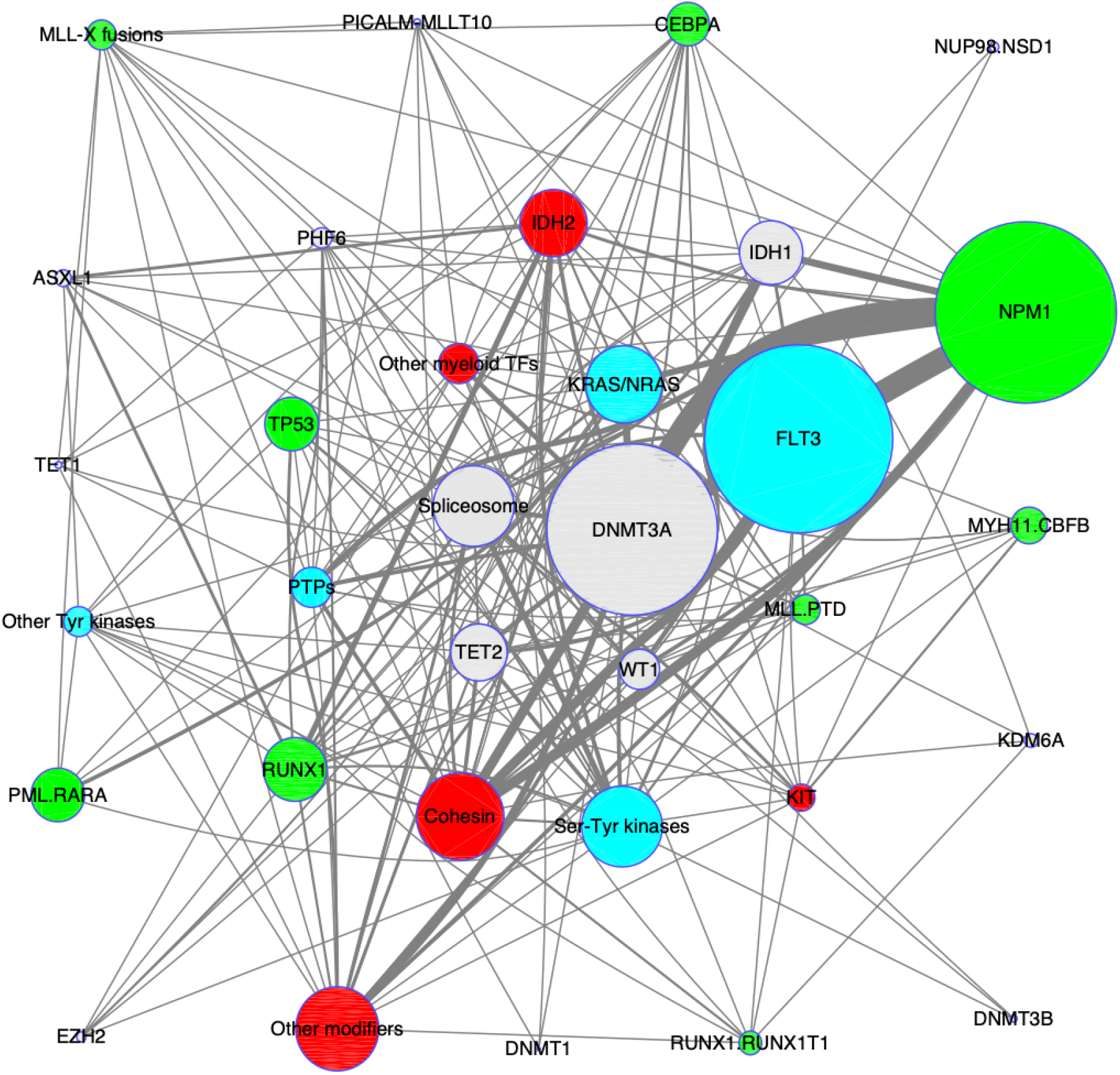
Graphical representation of AML data and three pathways identified with *α* = 0.70.

If we extract three more pathways, the results are as follows:

- Pathway4 = ‘PHF6’, ‘DNMT3A’, ‘KIT’, coverage = 65, coverage/gene = 21.67, indep = 4, measure = 5.42
- Pathway5 = ‘WT1’, ‘TET2’, ‘IDH2’, ‘PTPs’, coverage = 61, coverage/gene = 15.25, indep = 4, measure = 3.81
- Pathway6 = ‘IDH1’, ‘CEBPA’, ‘Spliceosome’, coverage = 56, coverage/gene = 18.67, indep = 4, measure = 4.67

All six pathways are depicted in Figure 4. Although the extra pathways were not reported by (Cancer Genome Atlas Research Network, 2013), our results suggest that these candidate altered cancer pathways may be worthy of review by disease experts.

**Figure 4:**
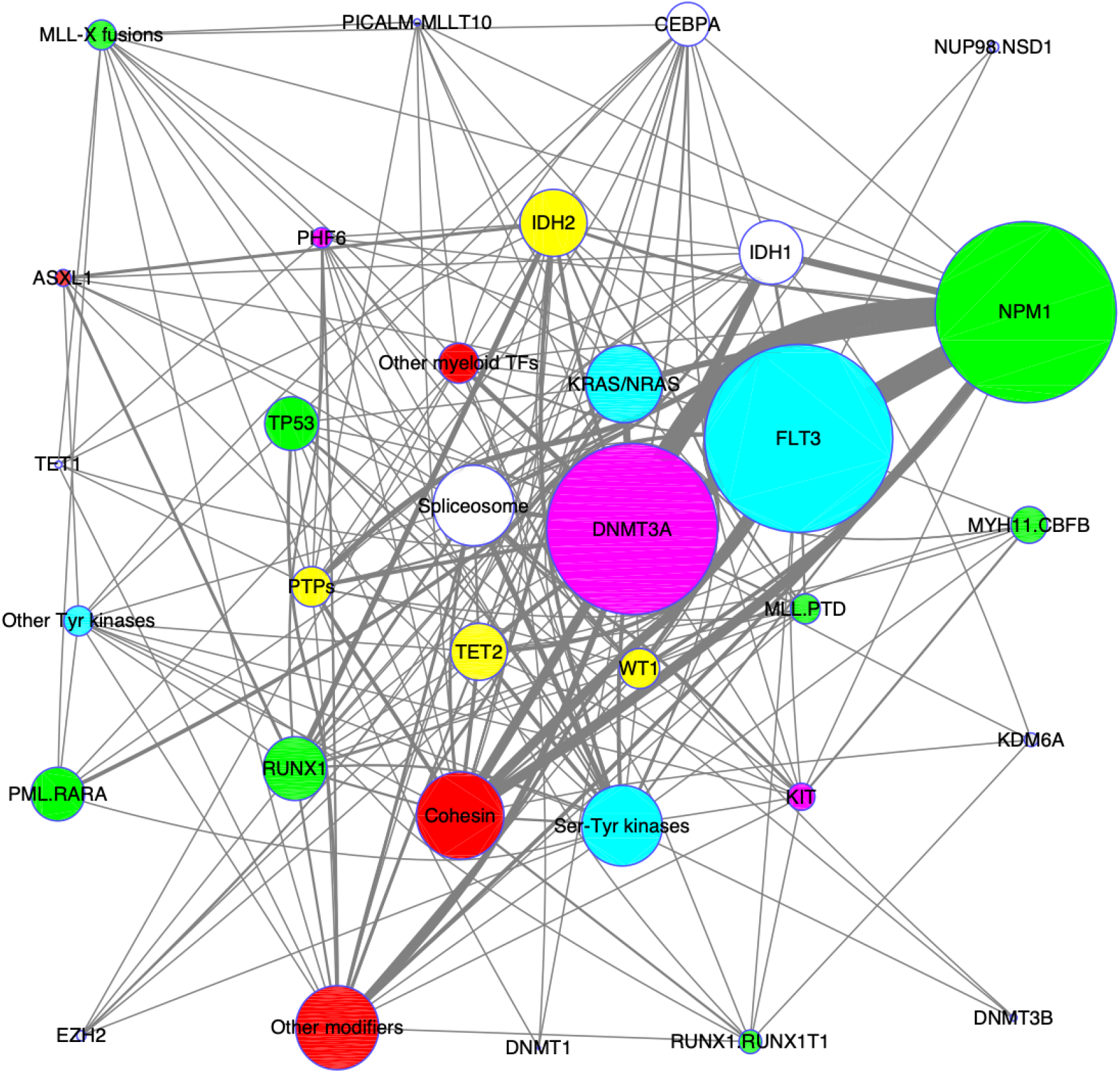
Graphical representation of AML data and six pathways identified with *α* = 0.45.

## A.2 GAMA Solver Results

In this section we present detailed results of pathways identified by GAMA.

## A.2.1 GAMA solver for three pathways (*k* = 3)

When *α* = 0.45, the results are as follows (solving time 1.68 sec):

- Pathway1 = ‘PML.RARA’ ‘MYH11.CBFB’ ‘RUNX1.RUNX1T1’ ‘NPM1’ ‘TP53’ ‘TET1’ ‘RUNX1’ ‘MLL.PTD’; coverage = 134, coverage/gene = 16.75, indep = 4, measure = 4.19
- Pathway2 = ‘DNMT1’ ‘IDH2’ ‘FLT3’ ‘Other Tyr kinases’ ‘MLL-X fusions’ ‘EZH2’ ‘KDM6A’; coverage = 102, coverage/gene = 14.57, indep = 4, measure = 3.64
- Pathway3 = ‘DNMT3B’ ‘Other myeloid TFs’ ‘NUP98.NSD1’ ‘ASXL1’ ‘Other modifiers’ ‘Cohesin’; coverage = 73, coverage/gene = 12.17, indep = 0, measure = Inf.

**Figure 5:**
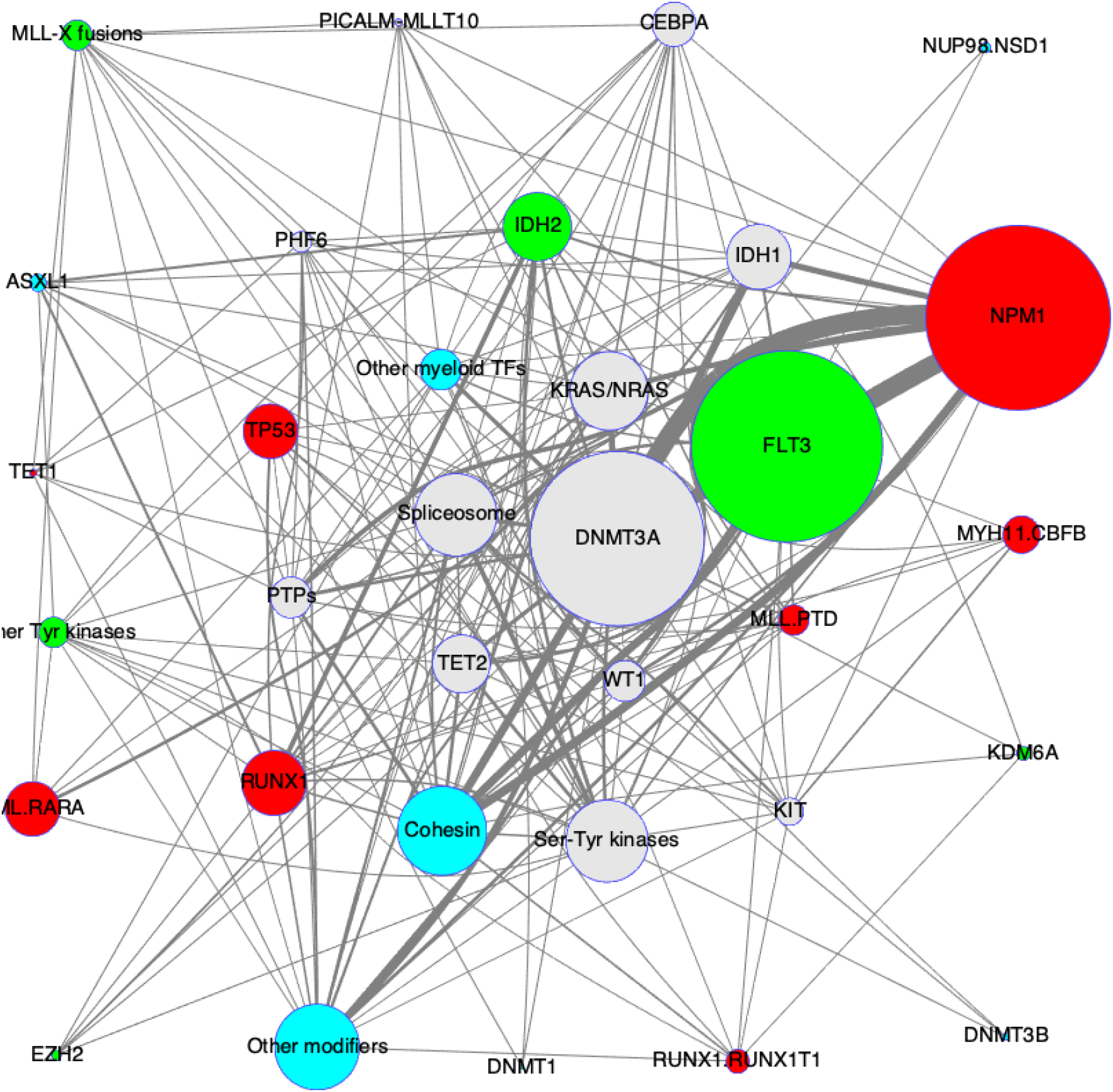
Graphical representation of AML data and three pathways identified with *α* = 0.45.

When *α* = 0.70, the results are as follows (solving time 1.71 sec):

- Pathway1 = ‘PML.RARA’ ‘MYH11.CBFB’ ‘RUNX1.RUNX1T1’ ‘NPM1’ ‘TP53’ ‘DNMT1’ ‘TET1’ ‘RUNX1’ ‘CEBPA’ ‘MLL.PTD’ ‘NUP98.NSD1’; coverage = 151, coverage/gene = 13.73, indep = 18, measure = 0.80
- Pathway2 = ‘IDH2’ ‘FLT3’ ‘Other Tyr kinases’ ‘PTPs’ ‘MLL-X fusions’ ‘EZH2’ ‘KDM6A’ ‘Spliceosome’; coverage = 137, coverage/gene = 17.13, indep = 26, measure = 0.66
- Pathway3 = ‘DNMT3B’ ‘KIT’ ‘KRAS/NRAS’ ‘Other myeloid TFs’ ‘ASXL1’ ‘Other modifiers’ ‘Cohesin’; coverage = 101, coverage/gene = 14.43, indep = 18, measure = 0.80

**Figure 6:**
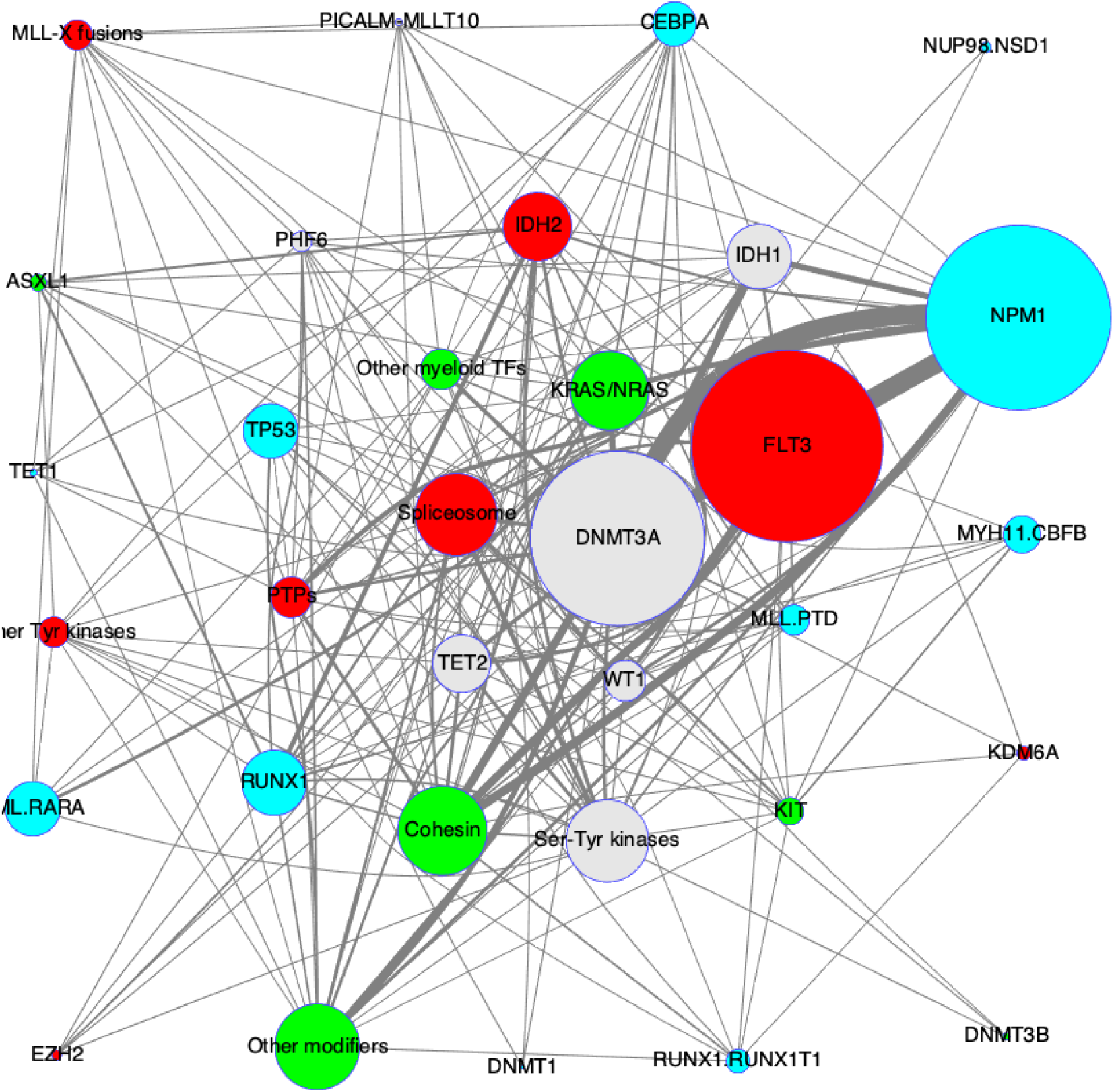
Graphical representation of AML data and three pathways identified with *α* = 0.45.

## A.2.2 GAMA solver six pathways (*k* = 6)

When *α* = 0.45, the results are as follows (solving time 10.04 sec):

- Pathway1 = ‘PICALM-MLLT10’ ‘PHF6’ ‘DNMT1’ ‘IDH1’ ‘CEBPA’ ‘MLL.PTD’ ‘Spliceosome’; coverage = 74, coverage/gene = 10.57, indep = 10, measure = 1.06
- Pathway2 = ‘MYH11.CBFB’ ‘RUNX1.RUNX1T1’ ‘NPM1’ ‘TP53’ ‘TET1’ ‘RUNX1’; coverage = 109, coverage/gene = 18.17, indep = 0, measure = Inf.
- Pathway3 = ‘WT1’ ‘DNMT3A’ ‘DNMT3B’ ‘Other Tyr kinases’ ‘MLL-X fusions’; coverage = 83, coverage/gene = 16.60, indep = 4, measure = 4.15
- Pathway4 = ‘PML.RARA’ ‘TET2’ ‘IDH2’ ‘KIT’ ‘PTPs’ ‘KDM6A’; coverage = 77, coverage/gene = 12.83, indep = 2, measure = 6.42
- Pathway5 = ‘Other myeloid TFs’ ‘NUP98.NSD1’ ‘ASXL1’ ‘Other modifiers’ ‘Cohesin’; coverage = 71, coverage/gene = 14.20, indep = 0, measure = Inf.
- Pathway6 = ‘FLT3’ ‘Ser-Tyr kinases’ ‘KRAS/NRAS’; coverage = 103, coverage/gene = 34.33, indep = 12, measure = 2.86

**Figure 7:**
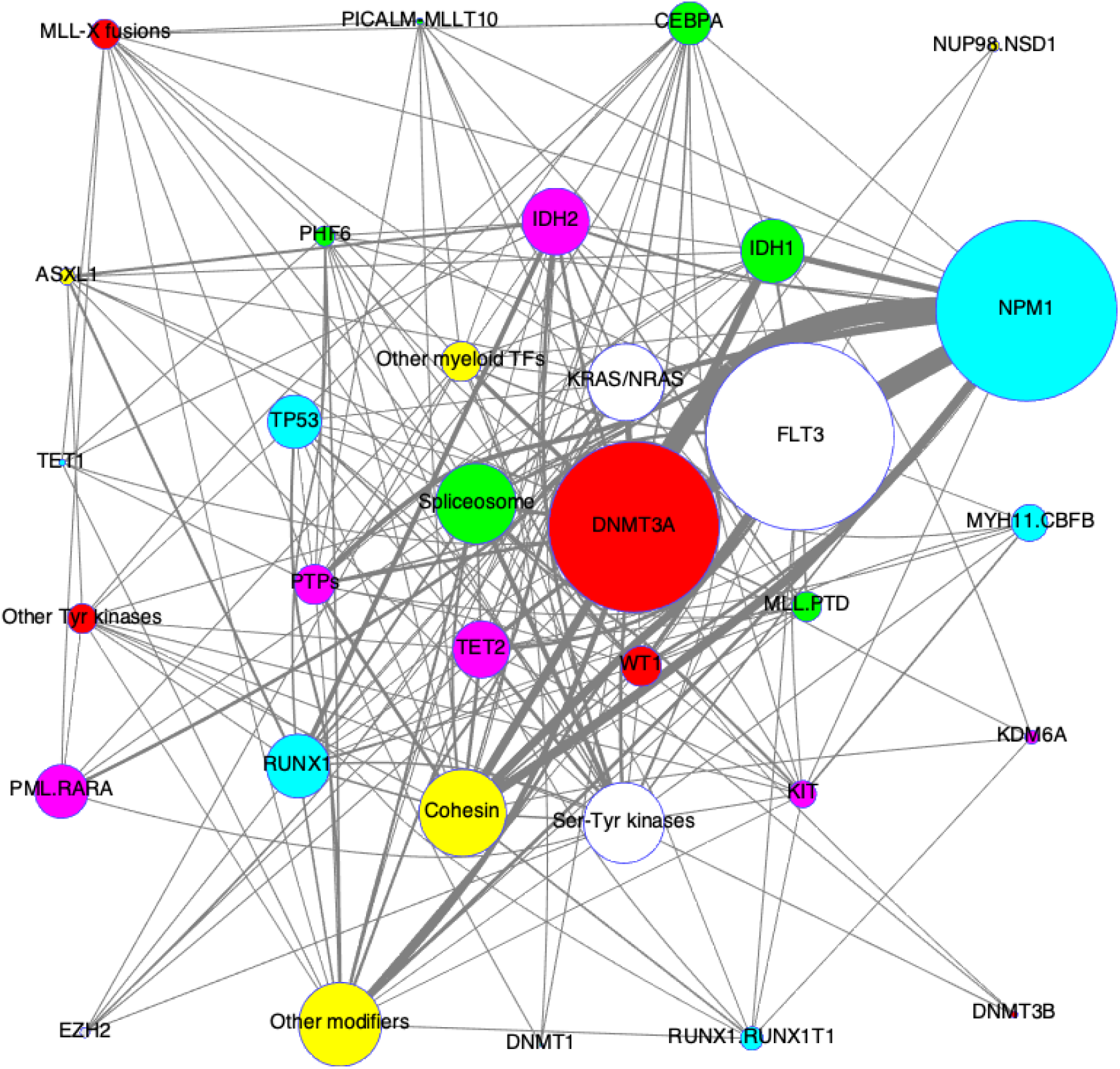
Graphical representation of AML data and six pathways identified with *α* = 0.45.

When *α* = 0.70, the results are as follows (solving time 10.48 sec):

- Pathway1 = ‘RUNX1.RUNX1T1’ ‘NPM1’ ‘TP53’ ‘TET1’ ‘RUNX1’; coverage = 98, coverage/gene = 19.60, indep = 0, measure = Inf.
- Pathway2 = ‘DNMT1’ ‘FLT3’ ‘CEBPA’ ‘EZH2’ ‘Spliceosome’; coverage = 97, coverage/gene = 19.40, indep = 10, measure = 1.94
- Pathway3 = ‘WT1’ ‘DNMT3A’ ‘DNMT3B’ ‘Other Tyr kinases’ ‘MLL-X fusions’; coverage = 83, coverage/gene = 16.60, indep = 4, measure = 4.15
- Pathway4 = ‘PML.RARA’ ‘MYH11.CBFB’ ‘PICALM-MLLT10’ ‘PHF6’ ‘TET2’ ‘IDH1’ ‘IDH2’; coverage = 91, coverage/gene = 13, indep = 8, measure = 1.63
- Pathway5 = ‘Other myeloid TFs’ ‘NUP98.NSD1’ ‘ASXL1’ ‘Other modifiers’ ‘Cohesin’; coverage = 71, coverage/gene = 14.20, indep = 0, measure = Inf.
- Pathway6 = ‘KIT’ ‘Ser-Tyr kinases’ ‘KRAS/NRAS’ ‘PTPs’ ‘MLL.PTD’ ‘KDM6A’; coverage = 80, coverage/gene = 13.33, indep = 12, measure = 1.67

**Figure 8:**
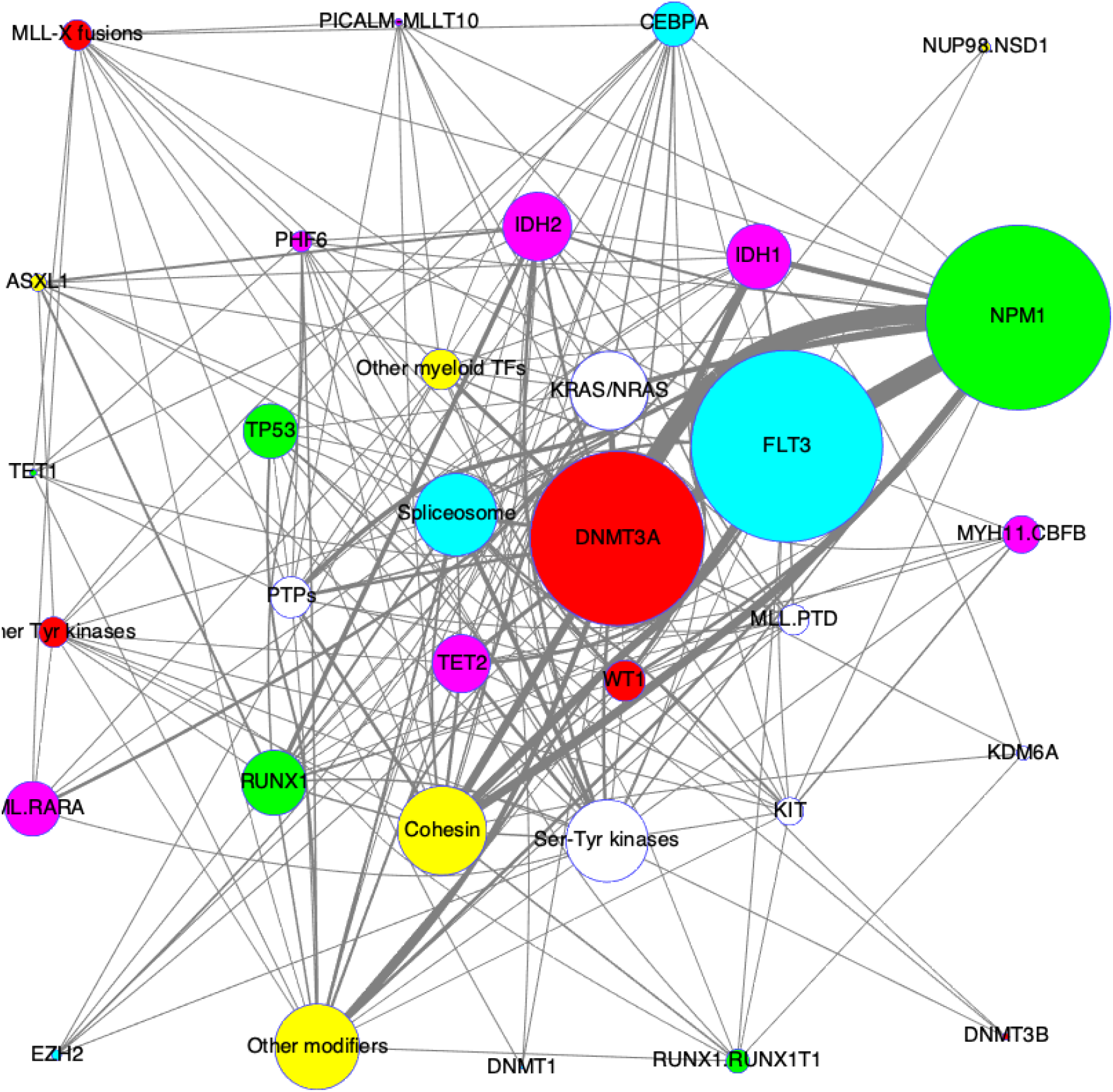
Graphical representation of AML data and six pathways identified with *α* = 0.70.

## B Graver Basis Optimization and the GAMA Solver

This section contains a brief introduction to the Graver basis based integer optimization and the GAMA (Graver Augmented Multi-seed Algorithm) solver that we used for the altered pathway discovery problem. For details, interested readers should refer to (Alghassi et al., 2019a).

## B.1 Graver Basis Optimization

Let *f* : ℝ^*n*^ → ℝ be a real-valued function. We want to solve the general non-linear integer optimization problem

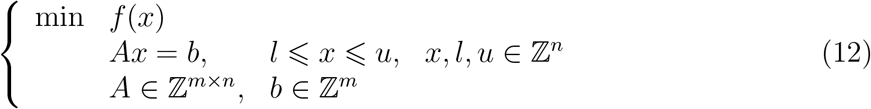

One approach to solving such a problem is to use an augmentation procedure: start from an initial feasible solution (which itself can be difficult to find) and take improvement steps (*augmentation*) until we reach the optimal solution. Augmentation procedures such as these need optimality certificates (or test sets). At every step in the optimization: either we can identify direction(s) that let us step towards better solution(s), or, if checking all possible directions offers no better solution, then we declare the optimality of the current solution. Note that it does not matter which feasible solution one begins from, nor the sequence of improving steps taken; the final stop is an optimal solution.

### Definition 1.

Let *x, y* ∈ ℝ^*n*^. We say *x* is *conformal* to *y*, written *x* ⊑ *y*, when *x*_*i*_*y*_*i*_ ≥ 0 (*x* and *y* lie on the same orthant) and |*x_i_*| ≤ |*y_i_*| for *i* = 1, …, *n*.

Suppose *A* is a matrix in ℤ^*m*×*n*^. Define lattice kernel of *A*:

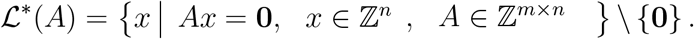

The notion of the Graver basis was first introduced in (Graver, 1975) for integer linear programs:

### Definition 2.

The Graver basis of integer matrix *A* is defined to be the finite set of minimal elements (*indecomposable* elements) in the lattice 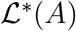. We denote by 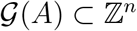 the Graver basis of *A*.

## B.1.1 Applicability of Graver bases as optimality certificates

Beyond integer linear programs (ILP) with a fixed integer matrix, Graver bases as optimality certificates have now been generalized to include several nonlinear objective functions (Onn, 2010):

- Separable convex minimization: min 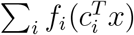 with *f_i_* convex.
- Convex integer maximization (weighted): max *f*(**W***x*), **W** ∈ ℤ^*d*×*n*^ with *f* convex on ℤ^*d*^.
- Norm *p* (nearest to *x*_0_) minimization: min ‖*x* − *x*_0_‖_*p*_.
- Quadratic minimization: min *x^T^ V x* where *V* lies in the dual of quadratic Graver cone of *A*
- Polynomial minimization: min *P* (*x*) where *P* is a polynomial of degree *d* that lies on cone *K_d_*(*A*), the dual of *d^th^* degree Graver cone of *A*.

It has been shown that only polynomially many augmentation steps are needed to solve such optimization problems (Onn, 2010).

## B.2 The GAMA Solver

The classical version of the Graver Augmented Multi-seed Algorithm (GAMA) solver primarily addresses hard-to-solve, practical, non-linear integer programming problems that have a specially structured linear constraint matrix. An important subset of these classes—including Cardinality Boolean Quadratic Problems (CBQP), Quadratic Semi-Assignment Problems (QSAP), and Quadratic Assignment Problems (QAP)—have two features: (1) their Graver bases can be calculated systematically, and (2) multiple feasible solutions that are uniformly spread out in the space of solutions can be likewise systematically constructed. The hardness of such problems, then, stems from the non-convexity of their nonlinear cost function, and not from the Graver basis or the ability to find feasible solutions.

An essential concept in test set (optimality certificate) based optimization in general, and Graver basis based optimization in particular, is separating the objective function from the constraints in the decomposition. It is known that, given the Graver basis of a problem’s integer matrix, any *convex* non-linear problem can be globally optimized with a polynomial number of moves—augmentations—from any arbitrary initial feasible solution (Onn, 2010). What is novel and different in GAMA is our recognition that for a wide class of hard problems, the matrix *A* has a special structure that allows us to obtain (a) Graver basis elements and (b) many feasible solutions that are spread out, using classical methods that are simple enough to be systematically algorithmized.

Suppose further that the *non-convex objective function* can be viewed as *many convex functions stitched together* (like a quilt). Thus, the entire feasible solution space can be seen as a collection of parallel subspaces, each with a convex objective function. If we have the Graver basis for the constraint matrix, and a feasible solution in every one of these subregions, then, putting this all together, an algorithm that can find the optimal solution is as follows:

- Find the Graver basis.
- Find a number of feasible solutions, spread out so that there is at least one feasible solution in each of the sub-regions that has a convex objective function.
- Augment along the Graver basis from each of the feasible solutions (‘seeds’) until you end up with a number of local optimal solutions (one for each seed).
- Choose the best from among these locally optimal solutions.

## B.2.1 QSAP and QAP problems

The Quadratic Semi Assignment problem formulation is as follows:

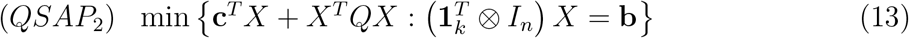

This form matches exactly with our formulation of (7), when only an orthogonality constraint is used.

The Quadratic Assignment problem formulation is as follows:

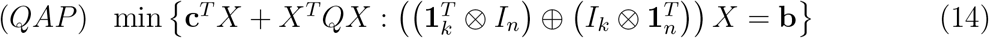

This form matches with our formulation of (7), when both orthogonality and cardinality constraints are used.

## C Quantum Annealing and the D-Wave Solver

## C.1 Optimization

Quantum annealing (Farhi et al., 2000; McGeoch, 2014) uses quantum adiabatic evolution (Kato, 1950) to solve optimization problems (QUBOs). This is done by slowly evolving the ground state of some known system into the sought ground state of the problem Hamiltonian. For instance, the D-Wave 2000Q (Harris et al., 2010; Johnson et al., 2011; Bunyk et al., 2014) implements quantum annealing using the time-dependent Hamiltonian

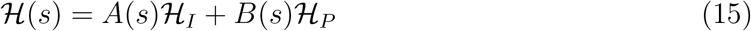

where *A*(*s*) is monotonically decreasing while *B*(*s*) is monotonically increasing with respect to normalized (slow) time 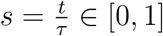. *τ* is the total annealing time. The initial Hamiltonian (with known ground state) is a transverse magnetic field 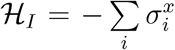 where 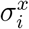 is the *i*^*th*^

Pauli *x*-matrix. The problem Hamiltonian 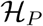 is (Barahona, 1982):

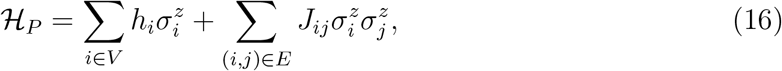

where the parameters *h_i_* and *J_ij_* encode the particular problem instance. The 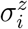 are Pauli *z*-matrices. A measurement of the final state (i.e. the end of the adiabatic evolution at *s* = 1) will yield a solution of the problem.

## C.2 D-Wave Solver

We had access to a D-Wave Quantum Annealer 2000Q™ (C16-VFYC solver) that is administered by the NASA Quantum Artificial Intelligence Laboratory (QuAIL). This processor operates at 20(±5)*milliKelvin* and was designed with 2048 qubits with a 95.5 qubits yield in its 16 by 16 block *Chimera* configuration. The *Chimera* structure has a limited connectivity, as shown in Figure 9. Therefore, the graph of a specific QUBO problem needs to be embedded into the graph of the hardware (Figure 9). Embedding an input graph into the hardware graph is a specific instance of a graph homomorphism, and its associated decision problem is in general NP-complete (Choi, 2008; Dridi et al., 2018).

**Figure 9:**
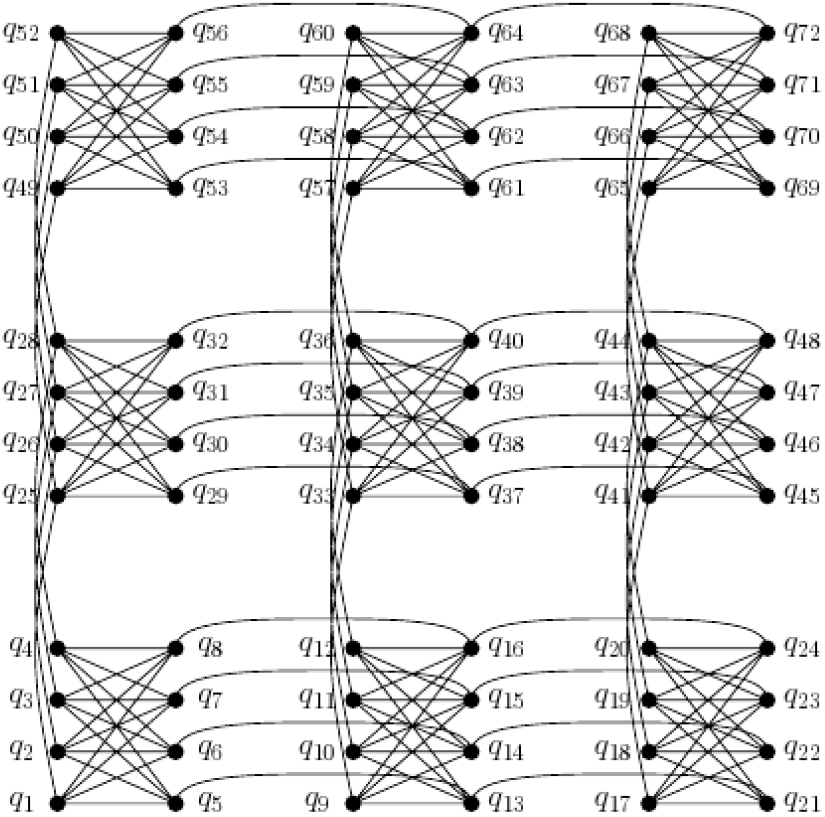
Schematic representation of a 3 by 3 block *Chimera*

## C.2.1 D-Wave SAPI

D-Wave’s software application programming interface (SAPI) is an application layer built to provide resource discovery, permissions, and scheduling for quantum annealing. The codes for this project are implemented using the Matlab SAPI (version 3.0). For embedding the problem graph into the hardware graph, and also for parameter settings, the sapiFindEmbedding (default parameters) and sapiEmbedProblem (adaptive problem per chain strength) software modules are used. To solve, sapiSolveIsing and sapiUnembedAnswer modules (with two distinct chainbreak strategies, *minimize energy* and *majority vote*) are used.

## Footnotes

1 This is also called representing graph, two section graph, clique graph, or Galifman graph.

2 **Q**_*card*_ = *I*_*k*_ ⊗ (*J*_*n*_ – 2*diag* (**s**)), where **s** = [*s*_1_ *s*_2_ … *s*_*k*_]^*T*^ is the cardinality vector.

3 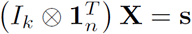 is the cardinality constraint in linear form, where **s** = [*s*_1_ *s*_2_ … *s*_*k*_]^*T*^ is the cardinality vector.

4 In the worst case each of m slack variables should be an integer covering [0, *n*].

## References

H. Alghassi. The algebraic QUBO design framework. Preprint, 2015. 6, 7

H. Alghassi, R. Dridi, and S. Tayur. GAMA: A Novel Algorithm for Non-Convex Integer Programs. arXiv:1907.10930 [quant-ph], July 2019a. URL http://arxiv.org/abs/1907. 10930. arXiv: 1907.10930. 3, 7, 9, 23

H. Alghassi, R. Dridi, and S. Tayur. Graver Bases via Quantum Annealing with Application to Non-Linear Integer Programs. arXiv:1902.04215 [quant-ph], Feb. 2019b. URL http://arxiv.org/abs/1902.04215. arXiv: 1902.04215. 3

F. Barahona. On the computational complexity of Ising spin glass models. Journal of Physics A: Mathematical and General, 15(10):3241, 1982. ISSN 0305-4470. doi: 10.1088/0305-4470/15/10/028. URL http://stacks.iop.org/0305-4470/15/i=10/a=028. 26

P. I. Bunyk, E. M. Hoskinson, M. W. Johnson, E. Tolkacheva, F. Altomare, A. J. Berkley, R. Harris, J. P. Hilton, T. Lanting, A. J. Przybysz, and J. Whittaker. Architectural Considerations in the Design of a Superconducting Quantum Annealing Processor. IEEE Transactions on Applied Superconductivity, 24(4):1–10, Aug. 2014. ISSN 1051-8223. doi: 10.1109/TASC.2014.2318294. 26

Cancer Genome Atlas Research Network. Genomic and epigenomic landscapes of adult de novo acute myeloid leukemia. The New England Journal of Medicine, 368(22):2059–2074, May 2013. ISSN 1533-4406. doi: 10.1056/NEJMoa1301689. 3, 9, 14, 17

F. Cheng, J. Zhao, and Z. Zhao. Advances in computational approaches for prioritizing driver mutations and significantly mutated genes in cancer genomes. Briefings in Bioinformatics, 17(4):642–656, 2016. ISSN 1477-4054. doi: 10.1093/bib/bbv068. 3

V. Choi. Minor-embedding in Adiabatic Quantum Computation: I. The Parameter Setting Problem. Quantum Information Processing, 7(5):193–209, Oct. 2008. ISSN 1570-0755. doi: 10.1007/s11128-008-0082-9. URL http://dx.doi.org/10.1007/s11128-008-0082-9. 26

G. Ciriello, E. Cerami, C. Sander, and N. Schultz. Mutual exclusivity analysis identifies oncogenic network modules. Genome Research, 22(2):398–406, Feb. 2012. ISSN 1088-9051. doi: 10.1101/gr.125567.111. URL http://www.ncbi.nlm.nih.gov/pmc/articles/PMC3266046/. 3

Y. Deng, S. Luo, C. Deng, T. Luo, W. Yin, H. Zhang, Y. Zhang, X. Zhang, Y. Lan, Y. Ping, Y. Xiao, and X. Li. Identifying mutual exclusivity across cancer genomes: computational approaches to discover genetic interaction and reveal tumor vulnerability. Briefings in Bioinformatics, 20(1):254–266, 2019. ISSN 1477-4054. doi: 10.1093/bib/bbx109. 3

C. M. Dimitrakopoulos and N. Beerenwinkel. Computational approaches for the identi-fication of cancer genes and pathways. Wiley Interdisciplinary Reviews: Systems Biology and Medicine, 9(1):e1364, 2017. ISSN 1939-005X. doi: 10.1002/wsbm.1364. URL https://onlinelibrary.wiley.com/doi/abs/10.1002/wsbm.1364. 3

R. Dridi, H. Alghassi, and S. Tayur. A Novel Algebraic Geometry Compiling Framework for Adiabatic Quantum Computations. arXiv:1810.01440 [quant-ph], Oct. 2018. URL http://arxiv.org/abs/1810.01440. arXiv: 1810.01440. 26

R. Dridi, H. Alghassi, and S. Tayur. The Topology of Mutated Driver Pathways. Preprint, CMU, 2019. 10

E. Farhi, J. Goldstone, S. Gutmann, and M. Sipser. Quantum Computation by Adiabatic Evolution. arXiv:quant-ph/0001106, Jan. 2000. URL http://arxiv.org/abs/quant-ph/0001106. arXiv: quant-ph/0001106. 26

J. E. Graver. On the foundations of linear and integer linear programming I. Mathematical Programming, 9(1):207–226, Dec. 1975. ISSN 0025-5610, 1436-4646. doi: 10.1007/BF01681344. URL https://link.springer.com/article/10.1007/BF01681344. 23

R. Harris, M. W. Johnson, T. Lanting, A. J. Berkley, J. Johansson, P. Bunyk, E. Tolkacheva, E. Ladizinsky, N. Ladizinsky, T. Oh, F. Cioata, I. Perminov, P. Spear, C. Enderud, C. Rich, S. Uchaikin, M. C. Thom, E. M. Chapple, J. Wang, B. Wilson, M. H. S. Amin, N. Dickson, K. Karimi, B. Macready, C. J. S. Truncik, and G. Rose. Experimental investigation of an eight-qubit unit cell in a superconducting optimization processor. Physical Review B, 82 (2):024511, July 2010. doi: 10.1103/PhysRevB.82.024511. URL https://link.aps.org/doi/10.1103/PhysRevB.82.024511. 26

M. W. Johnson, M. H. S. Amin, S. Gildert, T. Lanting, F. Hamze, N. Dickson, R. Har-ris, A. J. Berkley, J. Johansson, P. Bunyk, E. M. Chapple, C. Enderud, J. P. Hilton, K. Karimi, E. Ladizinsky, N. Ladizinsky, T. Oh, I. Perminov, C. Rich, M. C. Thom, E. Tolkacheva, C. J. S. Truncik, S. Uchaikin, J. Wang, B. Wilson, and G. Rose. Quan-tum annealing with manufactured spins. Nature, 473(7346):194–198, May 2011. ISSN 0028-0836. doi: 10.1038/nature10012. URL https://www.nature.com/nature/journal/v473/n7346/full/nature10012.html. 26

R. M. Karp. Reducibility among Combinatorial Problems. In R. E. Miller, J. W. Thatcher, and J. D. Bohlinger, editors, Complexity of Computer Computations, The IBM Research Symposia Series, pages 85–103. Springer US, 1972. ISBN 978-1-4684-2003-6978-1-4684-2001-2. doi: 10.1007/978-1-4684-2001-2_9. URL http://link.springer.com/chapter/10.1007/978-1-4684-2001-2_9. 3

T. Kato. On the Adiabatic Theorem of Quantum Mechanics. Journal of the Physical Society of Japan, 5(6):435–439, Nov. 1950. ISSN 0031-9015. doi: 10.1143/JPSJ.5.435. URL https://journals.jps.jp/doi/10.1143/JPSJ.5.435. 26

M. D. M. Leiserson, D. Blokh, R. Sharan, and B. J. Raphael. Simultaneous Identification of Multiple Driver Pathways in Cancer. PLOS Comput Biol, 9(5):e1003054, May 2013. ISSN 1553-7358. doi: 10.1371/journal.pcbi.1003054. URL http://journals.plos.org/ploscompbiol/article?id=10.1371/journal.pcbi.1003054. 3, 6, 8, 9

M. D. M. Leiserson, H.-T. Wu, F. Vandin, and B. J. Raphael. CoMEt: a statistical approach to identify combinations of mutually exclusive alterations in cancer. Genome Biology, 16: 160, 2015. ISSN 1474-760X. doi: 10.1186/s13059-015-0700-7. 3, 9

C. C. McGeoch. Adiabatic Quantum Computation and Quantum Annealing: Theory and Practice. Synthesis Lectures on Quantum Computing, 5(2):1–93, July 2014. ISSN 1945-9726. doi: 10.2200/S00585ED1V01Y201407QMC008. URL http://www.morganclaypool.com.proxy.lib.sfu.ca/doi/abs/10.2200/S00585ED1V01Y201407QMC008. 26

S. Onn. Nonlinear Discrete Optimization: An Algorithmic Theory. European Mathematical Soc., 2010. ISBN 978-3-03719-593-2. Google-Books-ID: kcD5DAEACAAJ. 23, 24

J. Preskill. Quantum Computing in the NISQ era and beyond. arXiv:1801.00862 [cond-mat, physics:quant-ph], Jan. 2018. URL http://arxiv.org/abs/1801.00862. arXiv: 1801.00862. 3, 10

E. Szczurek and N. Beerenwinkel. Modeling Mutual Exclusivity of Cancer Mutations. PLOS Comput Biol, 10(3):e1003503, Mar. 2014. ISSN 1553-7358. doi: 10.1371/journal.pcbi.1003503. URL http://journals.plos.org/ploscompbiol/article?id=10.1371/journal.pcbi.1003503. 3

F. Vandin, E. Upfal, and B. J. Raphael. De novo discovery of mutated driver pathways in cancer. Genome Research, 22(2):375–385, Feb. 2012. ISSN 1549-5469. doi: 10.1101/gr.120477.111. 3, 6, 7, 8, 9

Wikipedia. Hypergraph, 2019. URL https://en.wikipedia.org/w/index.php?title=Hypergraph&oldid=895361305. https://en.wikipedia.org/wiki/Hypergraph. 4

J. Zhao, S. Zhang, L.-Y. Wu, and X.-S. Zhang. Efficient methods for identifying mutated driver pathways in cancer. Bioinformatics (Oxford, England), 28(22):2940–2947, Nov. 2012. ISSN 1367-4811. doi: 10.1093/bioinformatics/bts564. 3

